# Optimiser Repairs for Living-Cell Self-Driving Laboratories: Performance and Redundancy under Experimental Constraints

**DOI:** 10.64898/2026.11.03.763926

**Authors:** Nura Brimo, Han Zhang, Dilek Çökeliler Serdaroğlu

## Abstract

Living-cell self-driving laboratories must optimise experiments despite delayed readouts, changing responses and limited sample availability. In simulations, we evaluate when optimiser repairs improve closed-loop performance and when platform constraints make them redundant, separating consumption of a sample from permanent loss of the ability to repeat its condition. We prove that exclusions and confirmation requests produce exactly the baseline trajectory when they act only on conditions already forbidden by the platform, share the same posterior and selection procedure, and release infeasible requested slots. This result covers the evaluated pending-exclusion, previous-batch and confirmation rules under permanent condition retirement; it does not cover arbitrary selection interventions. We then conduct 33,900 simulated policy runs with 200–300 paired campaign seeds per setting. A public PBMC reference supports characterisation of the target-label readout, while condition-response landscapes remain synthetic. Across 27 landscape, assay-size and kernel settings, previous-batch exclusion changes final normalised performance by −0.004 to +0.283, with positive differences surviving Holm adjustment in 13 settings. Effects are small on the rugged landscape. Permanent retirement can be harmful under noisy rugged responses. All three subsumed interventions reproduce every saved baseline event on 200 of 200 paired seeds. These results establish the importance of explicit availability constraints and evaluation across experimental regimes when assessing optimiser repairs.

## 1 Introduction

We ask whether repairs for delayed, changing or unreliable measurements improve closed-loop optimisation in living-cell laboratories, and when experimental constraints make those repairs redundant. An automated laboratory must decide both which experiment is informative and which experiment can actually be performed. These decisions interact. A planner may exclude an assay that is still running, request a repeat measurement, or prevent immediate reuse of a condition. A platform may already impose the same restriction through limited material or delayed sample preparation. Evaluating the additional rule without representing that platform constraint can therefore obscure what the optimiser changes.

Self-driving laboratories combine experimental planning, automated execution and model updates [11]. Living-cell experiments motivate particular care because the object measured, the preparation protocol and the underlying population are distinct. Sequencing consumes sampled cells [13]; it does not, by itself, make a culture condition impossible to prepare again. Likewise, a shortage of cells is not equivalent to assigning each condition a lifetime budget of one assay. A condition becomes nonrepeatable only when the experimental inventory and preparation process make a fresh evaluation unavailable within the campaign.

We examine the consequences of this distinction for finite-pool Bayesian optimisation. Our starting point is a simple identity: adding an exclusion already enforced by the platform cannot change the feasible set. If a requested replicate is also infeasible and its slot is returned to the ordinary selector, the complete decision can remain unchanged. We state the conditions under which this reasoning extends to exact equality of experimental trajectories. The result covers restrictions on pending or previously selected conditions when those conditions are unavailable. It does not cover all selection rules, and it does not imply that consuming a sample improves optimisation.

The empirical question is separate: when repetition is feasible, does preventing consecutive-batch repeats help? Batch local penalisation encourages separation within a batch [7], whereas replication can improve statistical estimation under noise [3]. There is consequently no general reason to expect an additional no-repeat rule to dominate. We test its sensitivity to response structure, assay sampling and the surrogate kernel, and compare it with permanent retirement and random search. We also evaluate pending-point handling, temporal adaptation and incumbent confirmation as distinct interventions.

We evaluate these interventions in computational experiments with synthetic condition-response functions. A public peripheral-blood mononuclear-cell (PBMC) reference is used to characterise the target-label readout; it does not supply measured responses to culture conditions. The study establishes sufficient conditions for exact intervention redundancy and quantifies performance changes across response structures, sampling noise levels and surrogate settings.

## 2 Experimental Framework

Let *X* ⊂ [0, 1]^6^ contain 1,024 candidate conditions. A campaign selects six distinct conditions per round for 20 rounds, giving 120 assay submissions. The same condition may occur in different rounds if the platform permits it. At the start of round *t, T*_*t*_ contains previously submitted conditions, *P*_*t*_ ⊆ *T*_*t*_ contains pending conditions, and *D*_*t*_ contains returned observations with their submission times. A delayed assay reports the state at submission, not the state at delivery.

The platform excludes *F*_*t*_ ⊆ *X*. We distinguish unlimited fresh preparations (*F*_*t*_ = ∅), a finite preparation budget, a resupply delay, and permanent condition retirement (*F*_*t*_ = *T*_*t*_). Figure 1 illustrates why the same optimiser rule can matter in one case and be redundant in another. These constraints describe preparation availability separately from the physical consequences of measurement.

**Figure 1:**
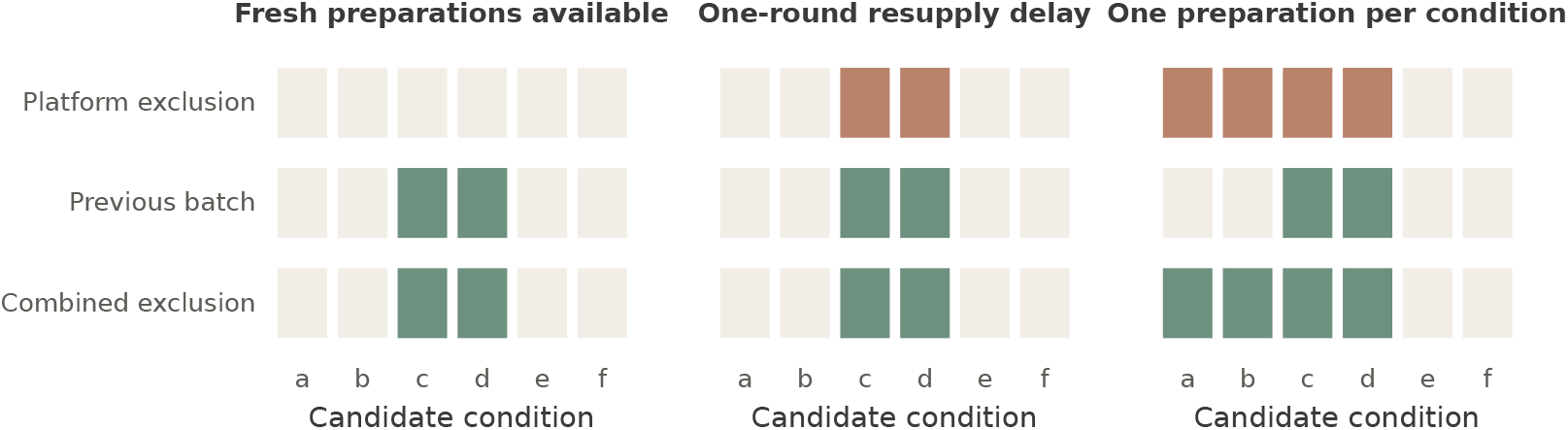
Platform constraints and additional exclusions. Conditions *a, b* were used earlier and *c, d* in the preceding batch. Filled cells indicate excluded conditions. A previous-batch rule adds restrictions when fresh preparations are available. It adds none during a one-round resupply delay or after permanent retirement. The diagram represents feasible candidate sets independently of policy performance.

An intervention is *model-level* if it changes the posterior for a fixed observation history and fixed settings. It is *selection-level* if it retains that posterior and changes only exclusions *G*_*t*_ or requested conditions *C*_*t*_. Each requested condition is checked against the platform constraints. An infeasible request is removed before allocating the remaining batch slots. All methods share this validation step. Confirmation changes future observations when it succeeds, but remains selection-level because it does not alter the model fitted to a given history.

The baseline uses a Matérn-5*/*2 Gaussian process and the acquisition

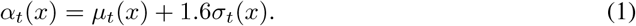

This is a fixed-weight upper-confidence heuristic inspired by GP-UCB [10]; we do not claim a regret guarantee for the chosen coefficient. We greedily assemble each batch, subtracting 2 exp[−∥*x* − *x*_*j*_∥ ^2^*/*(2*l*^2^)] after selecting *x*_*j*_, and mask exact within-batch duplicates. This fixed Gaussian repulsion heuristic differs from the Lipschitz-based penaliser of González et al. [7].

After observations due in a round have been incorporated, the recommendation is the observed condition with the largest posterior mean. We evaluate it after any immediate readouts from the current batch. If no result has returned, the score is zero. The primary outcome is

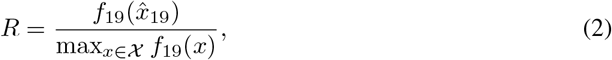

where rounds are indexed from 0. We also record distinct conditions sampled and whether a recommendation reaches 90% of the contemporaneous optimum at any round. Observations returned after round 19 are excluded from the final model. The denominator is always the maximum over the entire fixed candidate pool, including retired conditions; *R* measures identification of a good protocol, not the value of an executable next assay.

## 3 Theoretical Analysis of Intervention Redundancy

### Proposition 1

(Redundancy under Platform Constraints). *Consider two campaigns with the same initial state, posterior routine, acquisition, batch size, tie-breaking and event-indexed randomness. One additionally applies exclusions G*_*t*_ *and requests C*_*t*_. *Suppose G*_*t*_ ⊆ *F*_*t*_ *and C*_*t*_ ⊆ *F*_*t*_ *at every shared history. Infeasible requests are discarded without consuming batch slots or changing any other state. Then the campaigns select identical batches and produce identical observations, recommendations and derived performance indicators at every round*.

*Proof*. Assume the histories agree at the start of a round. The posterior and acquisition agree. Since *F*_*t*_ ∪ *G*_*t*_ = *F*_*t*_, both selectors have the same feasible set. Every requested condition is forbidden, so all six slots are available to the same batch selector. Identical tie-breaking produces identical batches.

The environment then returns identical observations and updates the same inventory, because selected events receive the same random inputs. Thus the histories agree in the next round. The common initial state completes the induction.

An exclusion already imposed by the platform leaves the feasible set unchanged. Similarly, an infeasible confirmation request leaves the selected batch unchanged when its reserved slot is released. Under permanent retirement, pending conditions, the previous batch and an observed incumbent all lie in *T*_*t*_ = *F*_*t*_. The three corresponding interventions satisfy the proposition. A resupply delay can already make previous-batch exclusion redundant without permanent retirement.

### Corollary 1

(Effective Intervention Sets). *For a fixed history, the additional exclusions are exactly G*_*t*_ \ *F*_*t*_, *and the feasible requests are C*_*t*_ \ *F*_*t*_. *Replacing an intervention by these residual sets leaves its action unchanged when request ordering and conflict handling are preserved*.

This is a set identity, not an upper bound on performance. Nonempty residual sets permit, but do not require, a different decision; they need not improve its value. For example, with candidate scores *α*(*a*) *> α*(*b*) *> α*(*c*) and *F* = {*c*}, an intervention *G* = {*a*} changes the selected condition from *a* to *b*. Thus exclusions of untested conditions, including cost or safety restrictions, are not covered merely because some conditions have been retired. Conversely, excluding only *b* would leave the selected condition unchanged. These examples also show why observing equal trajectories does not establish the proposition’s assumptions.

## 4 Methods

### 4.1 Biological Reference and Observation Model

The reference is the processed PBMC68k subset distributed with Scanpy 1.11.5: 700 cells, 765 genes and ten annotated types [12, 9, 13]. We use the first 20 bundled principal components and compute one centroid per annotated type. Thirteen cells are labelled CD34^+^ (1.86%). The unweighted mean within-type distance is 7.623, the mean pairwise centroid distance is 13.016, and their ratio is 0.586. The cell-weighted within-type distance is 7.094; the two averages should not be interchanged.

We assess the target-label readout within the reference dataset before simulating campaigns. In the reference itself, nearest-centroid assignment identifies all 13 CD34^+^ cells and assigns none of the 687 remaining cells to that class. This is a resubstitution check on a small reference, not validation on new donors. With no expression-space perturbation, independent empirical cell resampling therefore gives exactly the binary observation model

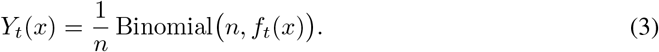

We use this equivalent, computationally efficient sampler. The expression geometry does not add classification error in these experiments. Cells per assay *n* ∈ {25, 200, 1000} control sampling noise; they are simulated cell draws, not independent biological replicates. Statistical replication is across campaigns, consistent with the distinction between within-experiment observations and independent experiments [8].

Three synthetic landscapes vary the relationship between conditions and target fraction: a Gaussian peak, the maximum of four weighted Gaussian peaks, and a random Fourier surface with shorter spatial variation. All are scaled to a maximum target fraction of 0.55 over the candidate pool; a 0.01 background is added before the remaining 0.54 amplitude. The scaling does not use observed outcomes and is hidden from the optimiser. The peak locations and Fourier coefficients are sampled afresh for each campaign. Appendix A gives the complete formulas. These functions test optimisation mechanisms; they do not represent measured PBMC responses to media, drugs or passage conditions.

### 4.2 Delayed Observations and Pending Evaluations

The latency sweep uses *L*∈ {0, 1, 3, 5} rounds. Readouts from a submission at *t* are available before selection at *t* + *L*, except that *L* = 0 readouts arrive after the current batch is submitted. Pending exclusion adds *G*_*t*_ = *P*_*t*_ while leaving the posterior unchanged. The model-level alternative uses posterior-mean fantasies at every pending condition, following the kriging-believer principle [6]. Its fantasy-noise variance is 0.20 in standardised response units, four times the observation nugget. Pending points are retained as separate submitted assays, including repeated conditions.

For fixed hyperparameters and target scaling, conditioning on a posterior-mean fantasy leaves the posterior mean unchanged and reduces covariance. We implement that conditional covariance update directly, including before the first real measurement arrives. Thus fantasies are not counted as completed observations and cannot themselves become recommendations. The covariance update is performed without restandardising the observed responses.

### 4.3 Temporal Variation and Adaptive Models

For the drift sweep, a common Gaussian random walk translates the single peak, with coordinate-wise step standard deviations 0.01, 0.03 or 0.05 per round. Centres are clipped to [0.05, 0.95]^6^, and the pool maximum is recomputed. These values are simulation settings, not passage rates estimated from biological data. Experimental evidence that cell-line variation can alter responses motivates the problem, but does not calibrate this random walk [2].

We compare the stationary baseline with two model-level interventions. The temporal kernel is

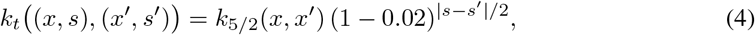

with the same temporal factor applied between observations and current predictions. This follows the covariance form studied by Bogunovic et al. [4]; our translated-peak process is not their assumed generative model. The sliding-window alternative retains only the most recent 32 returned observations, ordered by delivery and submission order. Recommendations remain eligible among all conditions with a returned observation.

### 4.4 Observation Corruption and Replicate Confirmation

Independently for each submitted assay, corruption replaces its target probability by a draw from Beta(1, 9) before binomial sampling. We vary corruption probability over 0, 0.05, 0.11, 0.20. This is a synthetic corrupted-readout model, not a mechanistic model of mycoplasma or a biological estimate of contamination frequency.

The confirmation arm considers the two observed conditions with the largest posterior means. It requests one incumbent replicate when their mean gap, divided by 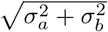, is below one. This score is a heuristic; it does not account for posterior covariance and is not a calibrated significance test. A feasible confirmation uses one of the six assay slots and counts toward the same budget as every other evaluation. An infeasible confirmation is discarded. The remaining slots use the baseline batch rule. This policy tests a specific replication heuristic rather than the general value of replication under noise [3].

### 4.5 Experimental Design and Statistical Analysis

The primary factorial study combines three landscapes, three assay sizes and three kernel length scales (*l* = 0.15, 0.30, 0.60). At each of 27 settings, 200 paired campaign seeds compare the baseline, previous-batch exclusion, permanent retirement and random search without replacement. The last two are controls: one tests a stronger availability restriction and the other measures the value of model-guided selection. The three repair sweeps use 300 paired seeds per setting, the single peak, *n* = 200 and *l* = 0.30. Additional availability experiments use 200 seeds for unlimited preparations, two preparations, one preparation, and resupply delays of one or three rounds. A separate 200-seed verification study combines *L* = 3, drift 0.03, corruption 0.11 and *n* = 25, with and without permanent retirement.

The primary contrast is the within-seed change in final performance. We report percentile 95% confidence intervals from 10,000 paired bootstrap resamples. Intervals are pointwise. Two-sided paired *t* tests receive Holm adjustment separately across the 27 primary robustness contrasts and the 18 repair contrasts; exploratory retirement contrasts form a separate family. We do not treat grid settings sharing seeds as independent replicates. Exact trajectory comparisons require identical submitted batches, delivered observation sequences, recommendations, scores and delivery counts. Success is computed from those same saved scores.

Hyperparameters were fixed for these comparisons without a performance-based search. Candidate pools, response surfaces and measurement uniforms are shared by paired arms; measurement randomness is indexed by campaign, round and condition. Different control-flow paths therefore cannot shift subsequent random draws. Distinct seed ranges are used for robustness, repairs, availability and structural verification. Appendix B records numerical settings and the event order.

## 5 Results

### 5.1 Effects of Previous Batch Exclusion

Previous-batch exclusion has a positive mean effect in 24 of 27 settings, but only 13 positive contrasts remain significant after Holm adjustment at 0.05 (Figure 2). At the default assay size and kernel, final performance increases from 0.810 to 0.953 on the single peak: paired change +0.143 (95% interval [0.098, 0.189]). On the four-peak surface it increases from 0.774 to 0.902, a change of +0.128 ([0.097, 0.160]). On the rugged surface the change is −0.002 ([−0.009, 0.007]). These comparisons support a regime-dependent effect, not a general failure of repeated evaluation.

**Figure 2:**
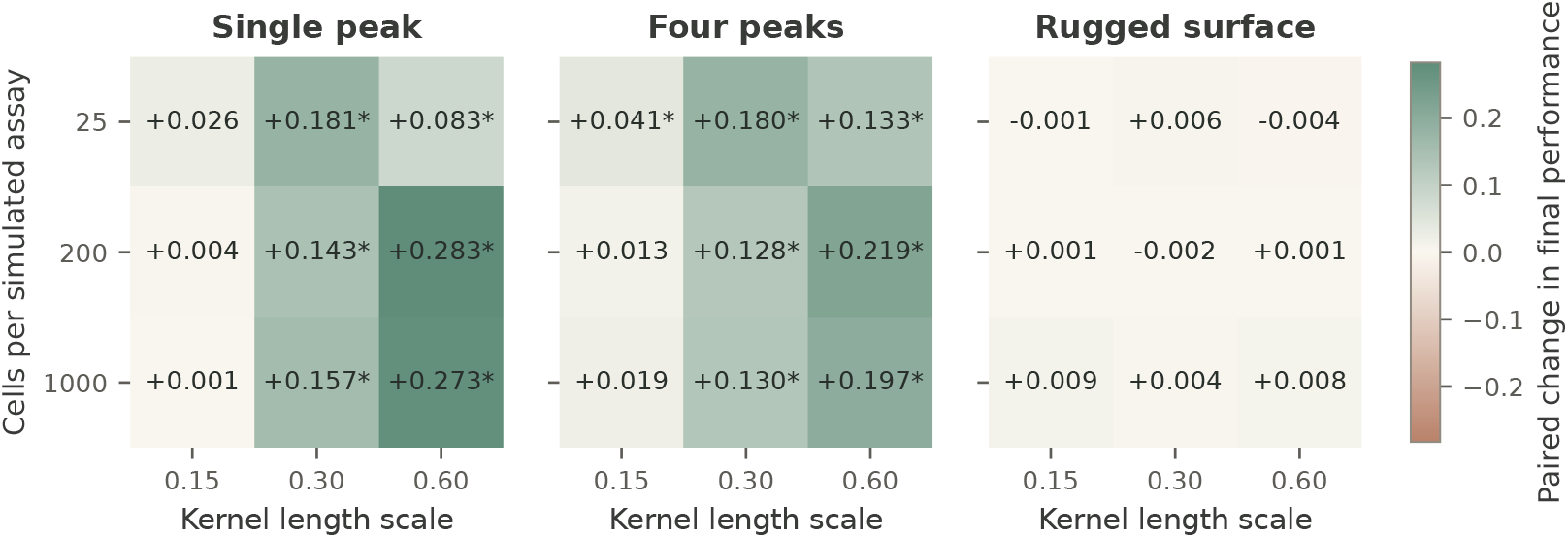
Sensitivity of previous-batch exclusion to response structure, assay size and kernel length scale. Each cell shows the mean paired change in final normalised performance over 200 seeds. Positive values favour the additional rule. Asterisks indicate *p*_Holm_ *<* 0.05 across all 27 comparisons. All three factors are varied jointly; the full paired intervals appear in Appendix C.

**Figure 3:**
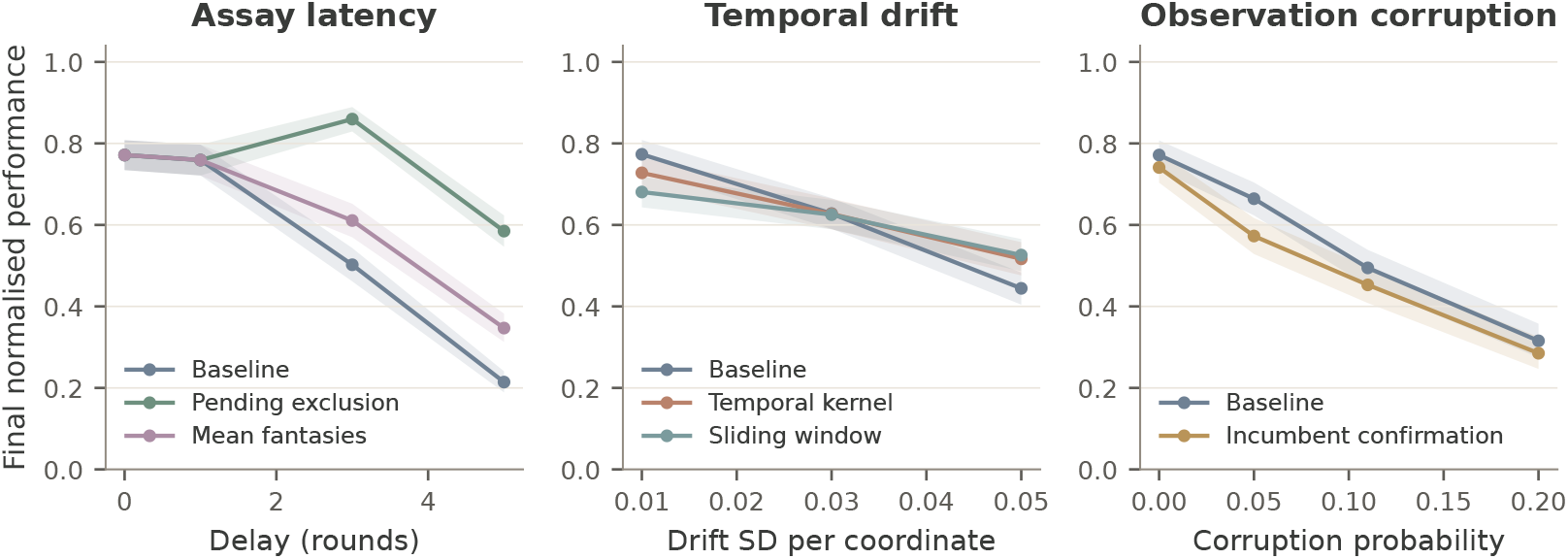
Performance under assay latency, temporal drift and observation corruption. Means and pointwise 95% bootstrap intervals over 300 paired seeds per setting. Each panel uses the single-peak landscape, 200 cells per assay and *l* = 0.30. The latency, drift and corruption sweeps are separate experiments. Mean fantasies improve the delayed loop in this implementation; temporal adaptation helps at the largest drift; the tested confirmation heuristic does not improve the mean at any corruption level. Full paired contrasts are in Table 2.

The largest primary gain occurs on the single peak with *n* = 200 and *l* = 0.60: +0.283 ([0.220, 0.346]). At *l* = 0.15 and the same assay size, the change is only +0.004 ([−0.002, 0.013]). Under the default settings, the previous-batch rule increases distinct conditions from 102.0 to 106.8 on the single peak, but more coverage is not sufficient for better performance. Random search samples 120 distinct conditions yet reaches 0.549, compared with 0.978 for model-guided permanent retirement (Appendix Figure 5).

Permanent retirement also supplies a counterexample to a universal diversification benefit. On the rugged landscape with 25 cells per assay, it reduces performance at all three length scales. The changes are −0.060, −0.039 and −0.069 for *l* = 0.15, 0.30, 0.60, respectively; all remain significant after the separate Holm adjustment for retirement contrasts. For the last setting the paired interval is [−0.089, −0.049]. Eliminating every repeat is therefore materially different from preventing only an immediate repeat.

### 5.2 Performance of Optimiser Repairs

At three rounds of latency, pending exclusion improves performance by +0.357 ([0.314, 0.401]), while mean fantasies improve it by +0.109 ([0.061, 0.156]). At five rounds, the corresponding changes are +0.371 and +0.133. All four remain significant after Holm adjustment. Under the stated delivery convention, zero- and one-round delays leave no pending condition at selection time, so both interventions coincide exactly with the baseline within those settings. Final performance differs between zero- and one-round latency because observations from the last batch remain pending in the latter setting.

At drift 0.05, the temporal kernel and sliding window improve performance by +0.072 ([0.027, 0.119]) and +0.082 ([0.029, 0.133]), with adjusted *p* = 0.027 for each. Neither intervention differs significantly from the baseline at drift 0.03 after multiplicity adjustment. At drift 0.01, the sliding window reduces performance by −0.093 ([−0.135, −0.049]). Confirmation decreases mean performance at every tested corruption level, although only the 0.05 setting remains significant after adjustment: −0.092 ([−0.131, −0.054]). It uses approximately 3.1–4.6 of the 120 assay slots on average. These results characterise the stated heuristics and do not rank all methods for temporal adaptation or robust replication.

### 5.3 Sample Availability and Trajectory Equivalence

With unlimited fresh preparations, the previous-batch rule improves performance by +0.195 ([0.148, 0.243]) on the independent availability seeds. With two preparations per condition its incremental effect is −0.004 ([−0.033, 0.025]), and all trajectories still differ. With one preparation, or a one- or three-round resupply delay, the rule is exactly redundant on all 200 seeds (Figure 4). The two-preparation setting therefore produces similar average performance despite different trajectories, whereas the more restrictive settings yield exact trajectory equivalence.

**Figure 4:**
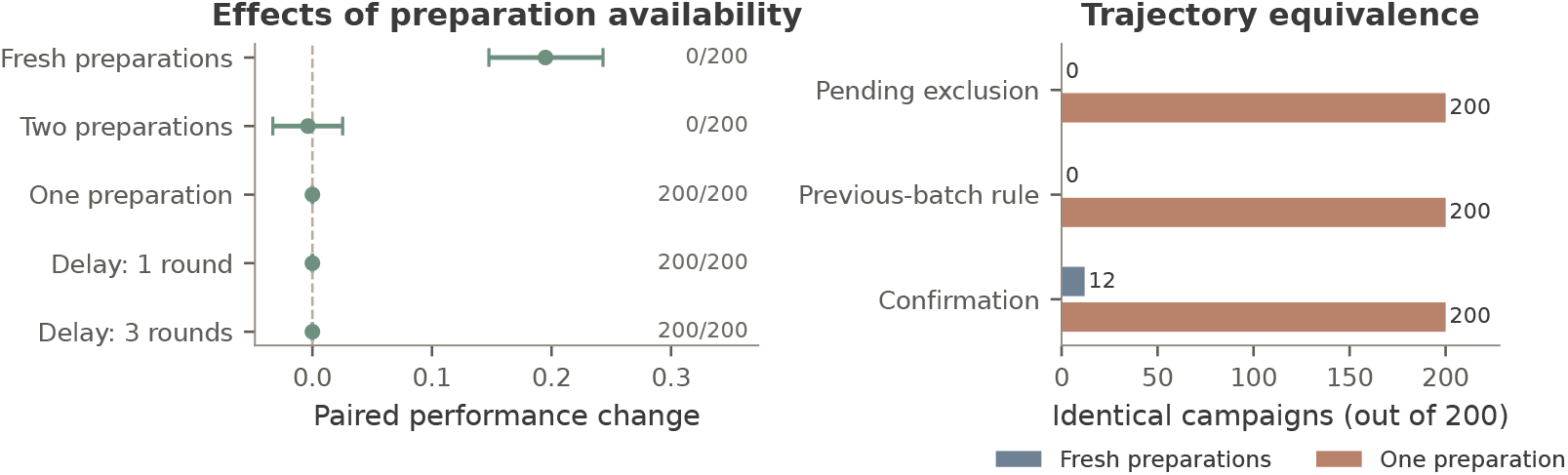
Effects of sample availability on performance and trajectory equivalence. Left: paired effect of previous-batch exclusion at five preparation regimes, with 95% intervals and counts of identical complete trajectories. Right: a separate verification study combines latency, drift, corruption and low assay size. Under permanent retirement, pending exclusion, previous-batch exclusion and confirmation each match all 200 baseline trajectories. With fresh preparations, equality is observed only in 12 confirmation campaigns.

**Figure 5:**
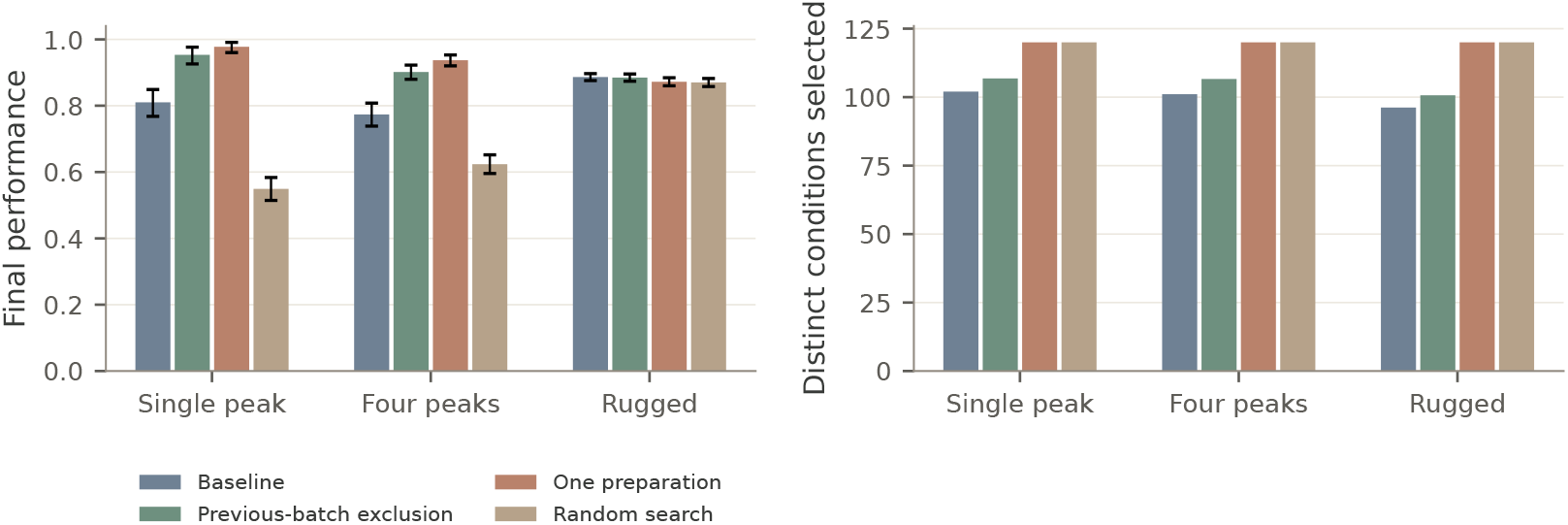
Recommendation performance and search coverage. The assay size is 200 cells and the kernel length scale is 0.30. Error bars in the left panel are pointwise 95% bootstrap intervals across 200 campaign seeds. The right panel shows mean distinct conditions selected from the 120-submission budget. Permanent retirement and random search without replacement both select 120 distinct conditions, so equal coverage need not imply equal recommendation quality.

In the combined-perturbation verification study, permanent retirement produces exactly the same batches, observations, recommendations and scores for the baseline and all three selection interventions on 200/200 seeds. Every arm has mean performance 0.226 and 34/200 successful campaigns.

Confirmation consumes zero replicate slots. Without retirement, pending exclusion and previous-batch exclusion match 0/200 baseline trajectories, while confirmation matches 12/200. The latter outcomes demonstrate possible divergence after removing the assumption; they are not a converse theorem. All 33,900 saved runs satisfy the specified batch and preparation constraints. Independent reconstruction verifies all 678,000 round-specific scores and their derived success indicators.

## 6 Discussion

The theoretical and empirical conclusions address different questions. Proposition 1 determines whether an intervention can change a decision under a specified state transition. It is independent of the response landscape, observation quality and optimiser accuracy. The robustness experiments determine whether a particular additional restriction improves the final recommendation when it can change the decision. Their answer depends on the tested regime.

For an automated laboratory, the immediate implication is to represent availability at the level of preparations and conditions. Consuming a well, waiting for a result and exhausting all preparations are different events. Before estimating an intervention’s mean benefit, one can test whether its residual exclusions or feasible requests are ever nonempty. When the proposition applies, identical trajectories are a useful implementation check. Trajectory equivalence requires agreement in submitted conditions, returned observations and recommendations; equality of rounded summary statistics is insufficient.

The experiments do not support a general prohibition on replication. On structured landscapes, preventing immediate repeats redirects the tested heuristic toward additional candidates. On the rugged landscape, those extra candidates offer little systematic benefit, and no tested grid setting establishes a reliable disadvantage for the previous-batch rule after multiplicity adjustment. The latter observation does not establish safety or equivalence: modest harms and other noise regimes remain possible. Planned replication can be valuable when uncertainty, rather than inadequate coverage, limits the decision [3].

The study has several limitations. The benchmark uses synthetic condition-response mappings, one small PBMC reference and a binary assay readout. It contains no newly collected cultures, donor validation, gene-regulatory dynamics, shared plate effects or biological dependence between simulated cell draws. The Gaussian likelihood is a convenience approximation to binomial observations, with a fixed standardised nugget. The study spans only six dimensions, 1,024 candidates and 120 submissions. Its explicit repulsion rule, fixed hyperparameters and simple confirmation threshold are not representative of all batch Bayesian optimisation methods; modern Monte Carlo acquisition frameworks offer substantially richer alternatives [1]. The 200-seed structural checks exercise the implementation, while the proof supplies the general conditional statement.

The practical cost of sample destruction also remains outside the performance score. Permanent retirement can increase search coverage while making a valuable condition unavailable for confirmation or subsequent use. Neither a high recommendation score nor zero software overhead establishes zero experimental cost. Similarly, changing to a less invasive readout, including image-based inference [5], should prompt a reassessment of availability and observation models; it does not automatically remove or create a diversification advantage.

## 7 Conclusions

Platform constraints can make additional optimiser rules exactly redundant, but only when those rules act entirely on conditions already excluded and do not otherwise alter the campaign. Permanent condition retirement is sufficient for the three selection interventions studied here; destructive measurement alone is not. When fresh preparations are available, the benefit of previous-batch exclusion depends on landscape structure and surrogate settings. Explicit availability models, paired comparisons and event-level verification provide a direct way to separate structural redundancy from an empirically weak intervention.

## A Synthetic Response Functions

For each seed, candidates are sampled independently and uniformly from [0, 1]^6^. Four centres *c*_*j*_ are sampled uniformly from [0.25, 0.75]^6^. The single-peak landscape uses

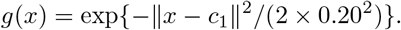

The four-peak landscape uses

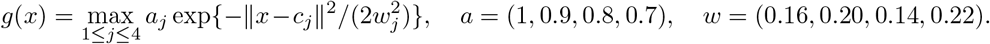

For the rugged landscape, *W* is a 6 64 matrix with independent *N* (0, 2^2^) entries and *ϕ*_*j*_ ~ Uniform(0, 2*π*). Define

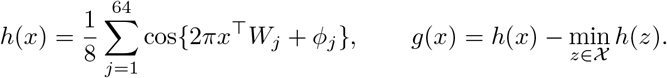

In every case *f* (*x*) = 0.01 + 0.54 *g*(*x*)*/* max_*z*∈*X*_ *g*(*z*). Different families have different distributions of near-optimal candidates even after this common amplitude scaling. Absolute scores should therefore not be used to rank their intrinsic difficulty. Within-family paired contrasts compare identical landscapes.

For drift, all centres receive the same accumulated Gaussian translation; only the first centre enters the single-peak surface. The translation at round *t* sums *t* independent increments, so round zero starts at the original centre. Coordinate clipping occurs after adding that accumulated translation. The landscape is normalised at each round. Corruption changes the readout probability, not the underlying objective. Repeated evaluations at different rounds receive independent measurement uniforms. Two policies evaluating the same condition in the same round receive the same uniform, whether their earlier actions agree or not.

## B Numerical Implementation

The spatial kernel has unit marginal variance and length scale *l*:

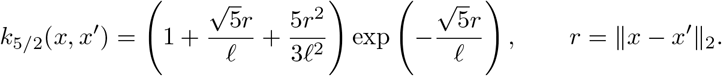

Observed responses are centred and divided by their population standard deviation, floored at 10^−6^. The standardised observation-noise variance is 0.05 and numerical diagonal jitter is 10^−8^. Both acquisition components are expressed on this standardised scale. Before the first observation the prior has zero mean and unit variance. Covariance systems are solved by Cholesky factorisation. A hard failure is raised if too few feasible candidates remain; none occurred in the reported runs. No hyperparameter optimisation or model-fit fallback is used.

Mean fantasies use the conditional covariance identity. If *K*^*D*^ denotes the covariance given real data and *P* the pending list, the adjusted covariance is

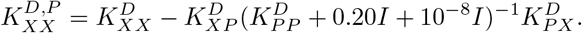

The posterior mean is unchanged because the imputed residual is zero. Fantasies are removed when their real observations arrive. The implementation verifies both this mean identity and the nonincrease of marginal variance. The final recommendation is obtained without fantasy conditioning, using the arm’s real-data posterior.

Each round follows five steps: (1) ingest previously submitted assays whose delivery time has arrived; (2) fit the surrogate and construct the platform mask; (3) validate any confirmation request and select the remaining feasible batch; (4) submit six assays and record each condition’s preparation count, immediately ingesting zero-delay readouts; and (5) refit with currently returned observations, record the recommendation and score, and retain pending events for subsequent rounds. Ties are broken by the first candidate index. Random search replaces the acquisition with independently generated candidate priorities and still respects the platform mask and within-batch uniqueness.

Under a preparation cap *m*, a condition is unavailable once its submission count reaches *m*. Under a resupply delay *d*, a condition last submitted at *s* is unavailable when *t* − *s* ≤ *d*. Thus a delay of one excludes exactly the preceding batch, while a delay of three excludes all submissions from the previous three rounds. These resupply rules concern new sample preparation and are separate from assay latency.

Seed ranges are 1000–1199 for the factorial study, 3000–3299 for repair sweeps, 5000–5199 for availability, and 7000–7199 for structural verification. Independent pseudorandom streams generate candidates, surface parameters, drift, assay uniforms, corruption indicators and replacement probabilities. The campaign seed is the independent inferential unit within each setting; policy arms sharing a seed form a paired comparison. Full run manifests record software versions and input fingerprints. The analysis used Python 3.12.14, NumPy 2.3.5 and SciPy 1.17.0.

## C Sensitivity Analysis of Previous Batch Exclusion

Table 1 reports all primary comparisons. A positive value favours previous-batch exclusion. Confidence intervals resample paired seed differences and are not adjusted for simultaneous coverage. The final column is the Holm-adjusted paired-test *p* value across the 27 rows. The accompanying CSV files provide complete performance and search-coverage summaries.

**Table 1:** Previous-batch exclusion minus baseline, 200 paired seeds per setting.

| Landscape | Cells | ℓ | Change | 95% interval | *p*<sub>Holm</sub> |
|---|---|---|---|---|---|
| single | 25 | 0.15 | +0.026 | [-0.012, +0.064] | 1 |
| single | 25 | 0.30 | +0.181 | [+0.115, +0.245] | 2.91e-06 |
| single | 25 | 0.60 | +0.083 | [+0.038, +0.130] | 0.00874 |
| single | 200 | 0.15 | +0.004 | [-0.002, +0.013] | 1 |
| single | 200 | 0.30 | +0.143 | [+0.098, +0.189] | 8.15e-08 |
| single | 200 | 0.60 | +0.283 | [+0.220, +0.346] | 1.96e-14 |
| single | 1000 | 0.15 | +0.001 | [-0.000, +0.002] | 1 |
| single | 1000 | 0.30 | +0.157 | [+0.118, +0.197] | 8.41e-12 |
| single | 1000 | 0.60 | +0.273 | [+0.219, +0.328] | 3.72e-17 |
| multimodal | 25 | 0.15 | +0.041 | [+0.020, +0.064] | 0.00421 |
| multimodal | 25 | 0.30 | +0.180 | [+0.128, +0.233] | 5.2e-09 |
| multimodal | 25 | 0.60 | +0.133 | [+0.088, +0.178] | 7.36e-07 |
| multimodal | 200 | 0.15 | +0.013 | [-0.002, +0.028] | 1 |
| multimodal | 200 | 0.30 | +0.128 | [+0.097, +0.160] | 3.9e-12 |
| multimodal | 200 | 0.60 | +0.219 | [+0.172, +0.268] | 4.18e-15 |
| multimodal | 1000 | 0.15 | +0.019 | [+0.006, +0.033] | 0.0919 |
| multimodal | 1000 | 0.30 | +0.130 | [+0.099, +0.161] | 1.94e-12 |
| multimodal | 1000 | 0.60 | +0.197 | [+0.152, +0.244] | 5.03e-13 |
| rugged | 25 | 0.15 | -0.001 | [-0.024, +0.020] | 1 |
| rugged | 25 | 0.30 | +0.006 | [-0.014, +0.025] | 1 |
| rugged | 25 | 0.60 | -0.004 | [-0.020, +0.011] | 1 |
| rugged | 200 | 0.15 | +0.001 | [-0.010, +0.012] | 1 |
| rugged | 200 | 0.30 | -0.002 | [-0.009, +0.007] | 1 |
| rugged | 200 | 0.60 | +0.001 | [-0.006, +0.009] | 1 |
| rugged | 1000 | 0.15 | +0.009 | [-0.001, +0.019] | 0.948 |
| rugged | 1000 | 0.30 | +0.004 | [-0.003, +0.011] | 1 |
| rugged | 1000 | 0.60 | +0.008 | [+0.002, +0.015] | 0.234 |

## D Comparative Performance of Optimiser Repairs

The repair sweeps share campaign seeds within each setting. Table 2 lists the paired contrasts and adjusted tests. A zero contrast is exact when no pending evaluation exists at selection time.

**Table 2:** Repair minus baseline, 300 paired seeds per setting. The 18 rows form one Holm-adjustment family.

| Factor | Value | Intervention | Change | 95% interval | *p*<sub>Holm</sub> |
|---|---|---|---|---|---|
| latency | 0 | Pending exclusion | +0.000 | [+0.000, +0.000] | 1 |
| latency | 0 | Mean fantasies | +0.000 | [+0.000, +0.000] | 1 |
| latency | 1 | Pending exclusion | +0.000 | [+0.000, +0.000] | 1 |
| latency | 1 | Mean fantasies | +0.000 | [+0.000, +0.000] | 1 |
| latency | 3 | Pending exclusion | +0.357 | [+0.314, +0.401] | 2.93e-40 |
| latency | 3 | Mean fantasies | +0.109 | [+0.061, +0.156] | 0.000172 |
| latency | 5 | Pending exclusion | +0.371 | [+0.329, +0.410] | 4.86e-47 |
| latency | 5 | Mean fantasies | +0.133 | [+0.097, +0.167] | 3.16e-11 |
| drift | 0.01 | Temporal kernel | -0.045 | [-0.090, -0.001] | 0.395 |
| drift | 0.01 | Sliding window | -0.093 | [-0.135, -0.049] | 0.000476 |
| drift | 0.03 | Temporal kernel | -0.001 | [-0.048, +0.045] | 1 |
| drift | 0.03 | Sliding window | -0.003 | [-0.050, +0.045] | 1 |
| drift | 0.05 | Temporal kernel | +0.072 | [+0.027, +0.119] | 0.0272 |
| drift | 0.05 | Sliding window | +0.082 | [+0.029, +0.133] | 0.0272 |
| corrupt | 0 | Incumbent confirmation | -0.031 | [-0.072, +0.011] | 1 |
| corrupt | 0.05 | Incumbent confirmation | -0.092 | [-0.131, -0.054] | 7.25e-05 |
| corrupt | 0.11 | Incumbent confirmation | -0.042 | [-0.080, -0.004] | 0.302 |
| corrupt | 0.2 | Incumbent confirmation | -0.030 | [-0.072, +0.010] | 1 |

## E Computational Verification

For each reported run, the verification script recomputes final performance and success from the stored score sequence, checks batch cardinality, within-batch uniqueness and preparation limits, and checks the previous-batch restriction whenever it applies. The structural test compares complete saved sequences, not merely rounded scores. With permanent retirement, every tested selection intervention matches the baseline on all 200 seeds and has the same 34 successful campaigns. The confirmation arm uses no confirmation slots. Without retirement, pending exclusion and previous-batch exclusion each differ on all 200 seeds; confirmation differs on 188 seeds and matches on 12. Divergence is an observed outcome in this implementation, not the converse of Proposition 1.

The reproducibility package includes the simulation code, reference embedding, calibration summary, per-round observations and submitted candidate indices. The PBMC data file has SHA-256 fingerprint `.e71d41e737c941559b7c57c9243bdb3d2c889c2adfdf00e3422ac6b46783676f`

## References

[1] Maximilian Balandat, Brian Karrer, Daniel R. Jiang, Samuel Daulton, Benjamin Letham, Andrew Gordon Wilson, and Eytan Bakshy. BoTorch: A framework for efficient monte-carlo bayesian optimization. In Advances in Neural Information Processing Systems, volume 33, pages 21524–21538, 2020. URL https://arxiv.org/abs/1910.06403.

[2] Uri Ben-David, Benjamin Siranosian, Gavin Ha, et al. Genetic and transcriptional evolution alters cancer cell line drug response. Nature, 560:325–330, 2018. doi: 10.1038/s41586-018-0409-3. URL https://doi.org/10.1038/s41586-018-0409-3.

[3] Mickaël Binois, Jiangeng Huang, Robert B. Gramacy, and Mike Ludkovski. Replication or exploration? sequential design for stochastic simulation experiments. Technometrics, 61(1): 7–23, 2019. doi: 10.1080/00401706.2018.1469433. URL https://arxiv.org/abs/1710.03206.

[4] Ilija Bogunovic, Jonathan Scarlett, and Volkan Cevher. Time-varying gaussian process bandit optimization. In Proceedings of the 19th International Conference on Artificial Intelligence and Statistics, volume 51 of Proceedings of Machine Learning Research, pages 314–323, 2016. URL https://proceedings.mlr.press/v51/bogunovic16.html.

[5] Eric M. Christiansen, Samuel J. Yang, D. Michael Ando, et al. In silico labeling: Predicting fluorescent labels in unlabeled images. Cell, 173(3):792–803.e19, 2018. doi: 10.1016/j.cell.2018.03.040. URL https://doi.org/10.1016/j.cell.2018.03.040.

[6] David Ginsbourger, Rodolphe Le Riche, and Laurent Carraro. Kriging is well-suited to parallelize optimization. In Computational Intelligence in Expensive Optimization Problems, pages 131–162. Springer, 2010. doi: 10.1007/978-3-642-10701-6_6. URL https://doi.org/10.1007/978-3-642-10701-6_6.

[7] Javier González, Zhenwen Dai, Philipp Hennig, and Neil Lawrence. Batch bayesian optimization via local penalization. In Proceedings of the 19th International Conference on Artificial Intelligence and Statistics, volume 51 of Proceedings of Machine Learning Research, pages 648–657, 2016. URL https://proceedings.mlr.press/v51/gonzalez16a.html.

[8] Samuel J. Lord, Katrina B. Velle, R. Dyche Mullins, and Lillian K. Fritz-Laylin. SuperPlots: Communicating reproducibility and variability in cell biology. Journal of Cell Biology, 219 (6):e202001064, 2020. doi: 10.1083/jcb.202001064. URL https://doi.org/10.1083/jcb.202001064.

[9] Scanpy developers. pbmc68k_reduced: Subsampled and processed 68k PBMCs, 2025. URL https://scanpy.readthedocs.io/en/latest/api/generated/scanpy.datasets.pbmc68k_reduced.html. Dataset distributed with Scanpy 1.11.5; accessed 29 October 2026.

[10] Niranjan Srinivas, Andreas Krause, Sham M. Kakade, and Matthias Seeger. Gaussian process optimization in the bandit setting: No regret and experimental design. In Proceedings of the 27th International Conference on Machine Learning, pages 1015–1022, 2010. URL https://arxiv.org/abs/0912.3995.

[11] Gary Tom, Stefan P. Schmid, Sterling G. Baird, et al. Self-driving laboratories for chemistry and materials science. Chemical Reviews, 124(16):9633–9732, 2024. doi: 10.1021/acs.chemrev.4c00055. URL https://doi.org/10.1021/acs.chemrev.4c00055.

[12] F. Alexander Wolf, Philipp Angerer, and Fabian J. Theis. SCANPY: Large-scale single-cell gene expression data analysis. Genome Biology, 19:15, 2018. doi: 10.1186/s13059-017-1382-0. URL https://doi.org/10.1186/s13059-017-1382-0.

[13] Grace X. Y. Zheng, Jessica M. Terry, Phillip Belgrader, et al. Massively parallel digital transcriptional profiling of single cells. Nature Communications, 8:14049, 2017. doi: 10.1038/ncomms14049. URL https://doi.org/10.1038/ncomms14049.

